# Lymphatic Malformations with Activating KRAS Mutations Impair Lymphatic Valve Development Through Matrix Metalloproteinases

**DOI:** 10.1101/2025.04.02.646922

**Authors:** Diandra M. Mastrogiacomo, Abbigail Price, Yuting Fu, Richa Banerjee, Luz A. Knauer, Kunyu Li, Ying Yang, George E. Davis, Michael T. Dellinger, Joshua P. Scallan

**Affiliations:** Department of Molecular Pharmacology and Physiology, University of South Florida, Tampa, FL, 33612; Department of Surgery, Department of Molecular Biology, University of Texas Southwestern Medical Center, Dallas, TX, 75390

**Author notes:** Corresponding author contact information: Joshua Scallan, Ph.D. Department of Molecular Pharmacology and Physiology, Morsani College of Medicine University of South Florida, 12901 Bruce B. Downs Blvd, MDC8 Tampa, FL 33612.

**Keywords:** lymphatic anomalies, lymphatic vessels, plasminogen activator pathway, shear stress, mechanotransduction

## Abstract

**BACKGROUND:** Lymphatic malformations (LMs) are lesions due to inherited or somatic mutations that lead to a defective lymphatic vasculature. Activating KRAS mutations have been identified recently in LM patients with lymphedema, chylous ascites, or life-threatening chylothorax. In a LM mouse model, KRAS mutations are associated with a loss of lymphatic valves, which has been proposed to cause chylothorax via retrograde lymph flow into the pleural space. However, the mechanisms underlying the loss of lymphatic valves are unknown.

**METHODS:** To investigate the mechanisms leading to valve loss, we combined the lymphatic-specific and tamoxifen-inducible *Flt4CreER*^*T2*^ with *Kras-loxP-stop-loxP-G12D* (*Kras*^*+/G12D*^) mice and *Prox1GFP* reporter mice to induce the restricted expression of KRAS-G12D and enable valve quantification in postnatal pups. Human dermal lymphatic endothelial cells (hdLECs) expressing KRAS-G12D were probed for changes in mRNA and protein expression with qRT-PCR, western blot, and gel zymography, and mechanistic studies were performed using 3D cell culture in collagen matrices.

**RESULTS:** Our data showed that lymphatic-specific expression of KRAS-G12D significantly attenuated valve development in the mesentery, diaphragm, and ear skin. qRT-PCR, western blot, and gel zymography using hdLECs expressing KRAS-G12D revealed the upregulation of the plasminogen activator (PA) pathway and matrix metalloproteinases (MMPs). The MMPs were sufficiently activated by plasmin, the product of the PA pathway, in hdLECs grown in a 3D collagen matrix, indicating a role for MMPs in the degradation of valve ECM core. Furthermore, a broad-spectrum MMP inhibitor given to *Flt4CreER*^*T2*^*;Kras*^*+/G12D*^ mice rescued lymphatic valve development.

**CONCLUSIONS:** We conclude that hyperactive KRAS signaling upregulates MMPs that become excessively activated by the upregulation of the PA pathway. MMPs then degrade the lymphatic valve ECM core preventing valve formation.

## INTRODUCTION

The lymphatic vasculature consists of a network of lymphatic vessels that absorb interstitial fluid and empty it into the bloodstream at the subclavian vein, a process required for fluid homeostasis.^1^ Fluid shear stress generated by lymph flow induces the growth of intraluminal bicuspid valves comprised of two layers of lymphatic endothelial cells (LECs) that encapsulate an extracellular matrix (ECM) core,^2-4^ and these valves are needed to prevent lymph reflux into tissues.^5^ The absence of lymphatic valves leads to a deadly condition known as chylothorax, where milky lymph leaks into the pleural space and airways, impairing respiration.^6^ Lymphatic valve defects were recently reported as a consequence of lymphatic malformations (LMs) caused by mutations in the *Kras* gene.^7,8^

LMs, also known as lymphatic anomalies, are congenital vascular lesions made up of dilated lymphatic vessels. Post-zygotic somatic gene mutations in the PI3K or MAPK pathways can lead to LMs, which impair lymphatic vessel function.^1,9^ LMs are commonly diagnosed at birth or within the first few years of life, but rarely in adults,^10^ with the overall prevalence as high as 1:4000 live births.^9^ LMs are classified into different types based on their pathology and consist of cystic lymphatic lesions, generalized lymphatic anomalies (GLA), central conducting lymphatic anomalies (CCLA), kaposiform lymphangiomatosis (KLA), and Gorham-Stout disease (GSD).^9^ Surgery or sclerotherapy treatment provides palliative care but is not curative, nor is it safe to perform in most patients as LMs are diffuse and usually form in vital anatomic locations, making complete surgical excision impossible.^11^ Most patients are given rapamycin, an mTOR inhibitor, as a first-line treatment in lieu of gene sequencing. However, many lesions do not respond to rapamycin, likely due to newly discovered mutations in genes in the RAS/MAPK pathway.^9,12^

Activating KRAS mutations were identified in the past few years in LM patients who present with lymphatic defects such as lymphangiectasia (pathological dilation of lymphatic vessels), chylous ascites, lymphedema, and/or chylothorax.^7,13,14^ A targeted cancer therapy called trametinib, a MEK inhibitor, was repurposed to treat LMs.^15^ Like rapamycin, trametinib can have adverse effects such as rash, dermatitis, diarrhea, and fatigue.^16^

Recently, the KRAS p.G12D activating mutation was linked to a severe loss of lymphatic valves in a LM mouse model.^7,8^ However, the mechanism whereby hyperactivated KRAS leads to valve loss is unknown. Lymphatic valve loss is associated with chylothorax because it allows retrograde lymph flow into the pleural space and lungs.^6^ Supporting this, LM patients with activating MAPK mutations can present with chylothorax, which increases disease mortality.^1,13^

Previous studies have demonstrated that hyperactivation of the MAPK pathway in cultured blood endothelial cells (BECs) and LECs expressing the same mutations identified in malformations (i.e. KRAS-G12V and ARAF-S214P, respectively) results in the loss of VE-cadherin from the cell membrane.^15,17^ We have demonstrated that VE-cadherin regulates mechanotransduction signaling in lymphatic vessels and is required for lymphatic valve formation and maintenance.^18^ Therefore, the current literature indicates that the loss of VE-cadherin from the cell membrane, and thus lack of mechanotransduction signaling, is a likely cause of valve loss in LMs.

Here, we show for the first time that the KRAS-G12D mutation in lymphatic vessels does not alter VE-cadherin expression or localization *in vivo* and thus cannot account for lymphatic valve defects in this type of LM. Instead, we demonstrate that hyperactivation of the KRAS/MAPK pathway leads to the upregulation of several components of the plasminogen activator (PA) pathway and matrix metalloproteinases (MMPs), resulting in the degradation of the ECM core of lymphatic valves, which impairs valve development.

## METHODS

For detailed experimental methods, materials, and statistical analyses, please see the Supplemental Material and Major Resource Table.

The following citations are for the detailed Materials and Methods and Major Resource Table sections.^19-25^

## RESULTS

### Expression of KRAS-G12D in mice results in lymphatic valve loss, chylothorax, and subsequent death by 8 weeks

To investigate putative mechanisms of valve loss in mice expressing the activating KRAS-G12D mutation, we bred the tamoxifen-inducible, lymphatic-restricted *Flt4CreER*^*T2*^ strain with *Kras-loxP-stop-loxP-G12D* mice (hereafter referred to as *Kras*^*+/G12D*^) to induce its expression in the majority of LECs.^19,26^ Tamoxifen (TM) was administered to newborn pups at postnatal day (P)0 and P2 to induce the expression of KRAS-G12D. The *Prox1GFP* reporter strain was used to visualize and quantify lymphatic valves (Figure 1A). First, we confirmed the high specificity and efficiency of the Cre activity at P7, P14 (data not shown), and P21 (Figure S1) using *Flt4CreER*^*T2*^*;Rosa26-mTmG* mice following the same TM schedule. Chylothorax was frequently observed in the P21 *Kras*^*+/G12D*^ mice, but was not observed in controls (Figure 1B). A survival analysis was performed with the *Kras*^*+/G12D*^ mice and every mouse that died was found to have chylothorax (17/17), with a median survival of 33 days (Figure 1C).

**Figure 1.**
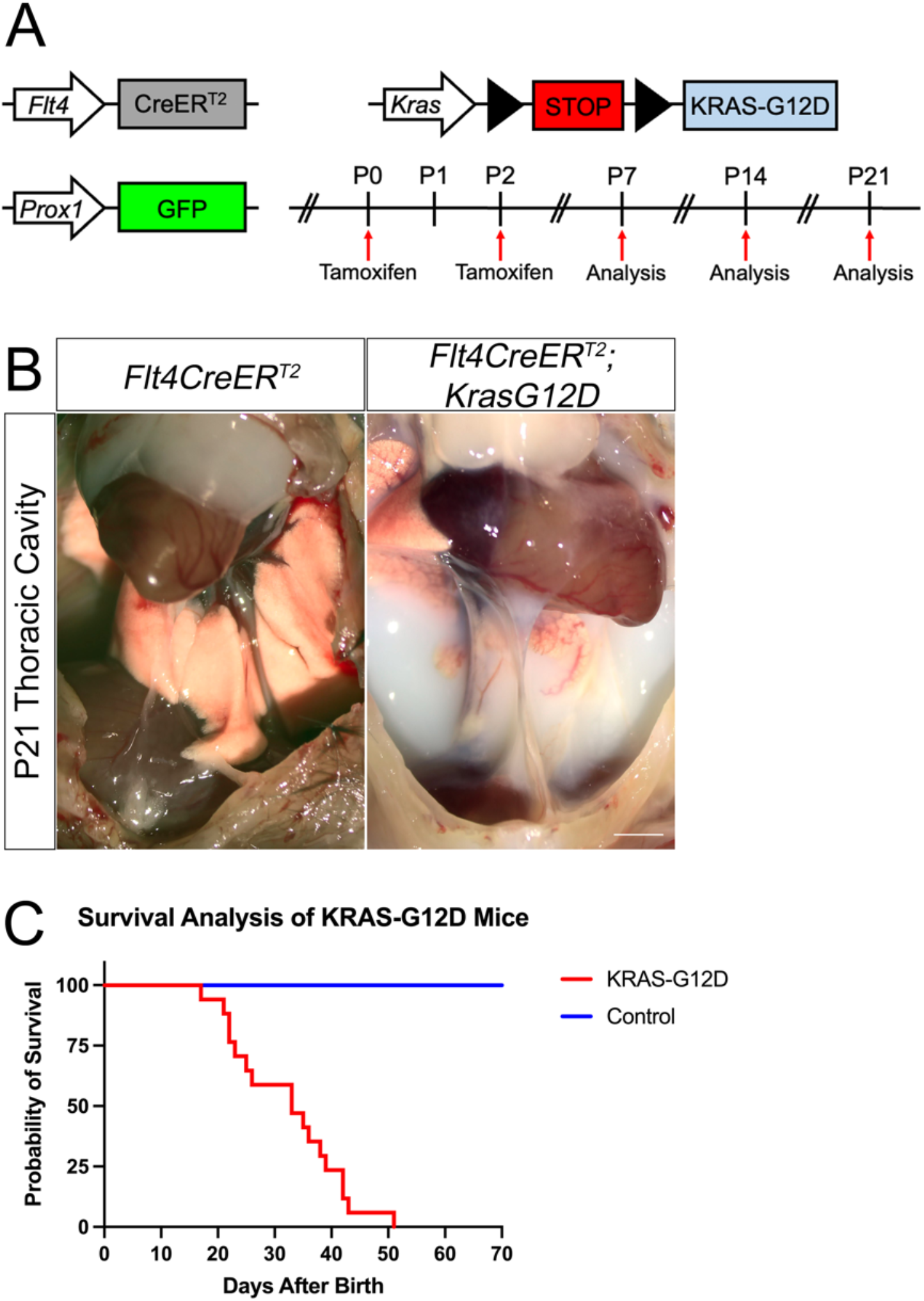
*Flt4CreER*^*T2*^*;Kras*^*+/G12D*^ do not live past 8 weeks due to chylothorax. **A**, The tamoxifen-inducible, lymphatic-restricted *Flt4CreER* ^*T2*^ was bred with *Kras-loxP-stop-loxP-G12D* mice to induce expression of KRAS-G12D in the majority of LECs. Tamoxifen (TM) was injected into newborn pups at postnatal day (P)0 and P2 to induce expression of KRAS-G12D. *Prox1GFP* reporter mice were used to visualize lymphatic vessels. Tissues were harvested at P7, P14 and/or P21 throughout this study. **B**, Chylothorax was observed in P21 *Flt4CreER*^*T2*^*;Kras*^*+/G12D*^ mice. Scale bar is 1 mm. **C**, Survival analysis was performed and each mouse that died was found to have chylothorax, with a median survival of 33 days. P<0.0001 by log-rank (Mantel-Cox) test, n=17 littermates per genotype.

Valves per millimeter (mm) of vessel length was quantified in the diaphragm and mesentery at P7, P14, and P21 in *Flt4CreER*^*T2*^*;Kras*^*+/G12D*^*;Prox1GFP* mice and *Flt4CreER*^*T2*^*;Prox1GFP* controls using NIH ImageJ (Figure 2). Valves were also quantified in the ear skin, but only at P21 when the ears were large enough to harvest and were fully developed. Lymphatic vessels from *Kras*^*+/G12D*^ mesenteries had 62%, 73%, and 78% fewer valves at P7, P14, and P21 than controls, respectively (Figure 2B, 2E, 2H). No significant differences were found in total vessel length in the mesentery at P7, P14 and P21 (Figure 2C, 2F, 2I).

**Figure 2.**
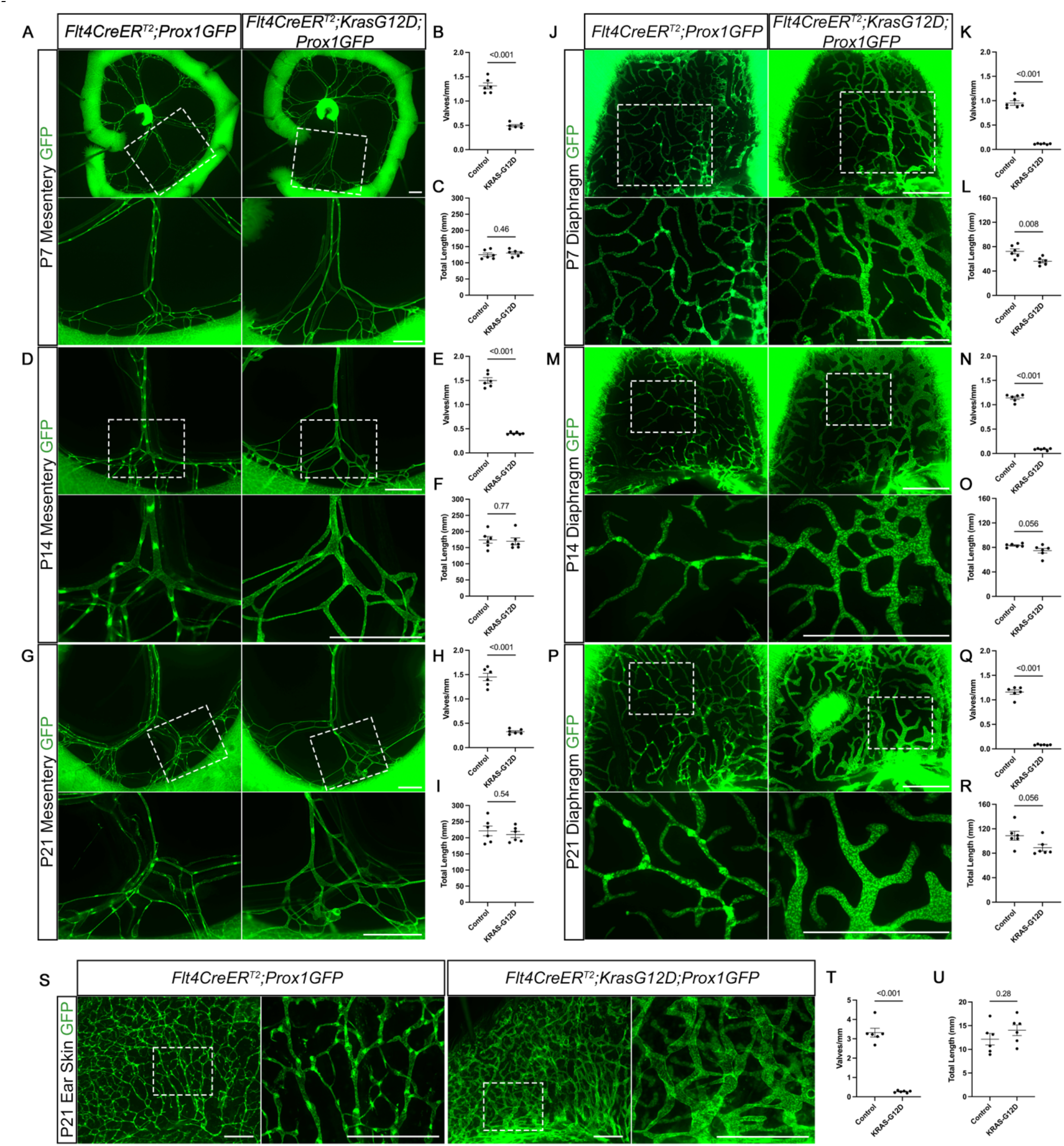
Lymphatic valve deficiency in *Flt4CreER*^*T2*^*;Kras*^*+/G12D*^ mice. Fluorescence images of the lymphatic vasculature represented by green fluorescent protein (GFP) in the mesentery (**A, D, G**), diaphragm (**J, M, P**) and ear skin (**S**) at postnatal day (P)7 (**A, J**), P14 (**D, M**) and P21 (**G, P, S**). Scale bar is 1 mm. **B, E, H, K, N, Q, T**, Valves per millimeter (mm) of vessel was quantified. **C, F, I, L, O, R, U**, Total length of vessels that was quantified. All values are mean ± SEM of n=6 per genotype. Analysis was performed with the unpaired Student *t* test.

In the diaphragm, *Kras*^*+/G12D*^ lymphatic vessels had 88%, 92%, and 93% fewer valves at P7, P14, and P21 compared to controls, respectively (Figure 2K, 2N, 2Q), while the total vessel lengths were significantly decreased by 22% only at P7, and no significant differences were found at P14 and P21 (Figure 2L, 2O, 2R). In the P21 ear skin, *Kras*^*+/G12D*^ lymphatic vessels had 92% fewer valves (Figure 2T) with no change in the total vessel length when compared to controls (Figure 2U).

### Many transcription factors regulating lymphatic valve development are downregulated upon expression of KRAS-G12D

Lymphatic valve development is regulated by transcription factors, such as FOXC2, GATA2, and PROX1.^27-29^ To assess the expression of these genes in KRAS-G12D expressing cells, we infected cultured human dermal LECs (hdLECs) with a lentivirus expressing the KRAS-G12D mutation, or an identical vector in which KRAS-G12D was replaced by mCherry as a control vector (CV). The hdLECs were then exposed to either static (no flow) or oscillatory shear stress (OSS) conditions for 48 hours (Figure S2). Quantitative real-time polymerase chain reaction (qRT-PCR) and western blot were performed for FOXC2, GATA2, and PROX1. Under both static and OSS conditions, *FOXC2, GATA2*, and *PROX1* were downregulated at the mRNA level (Figure S2A) and the protein level (Figure S2B) in hdLECs expressing KRAS-G12D.

To confirm these findings *in vivo*, we performed whole-mount immunostaining for the same transcription factors, along with VEGFR3, in P7 mesentery, P21 diaphragm, and P21 ear skin of *Kras*^*+/G12D*^ mice and controls (Figure S3). In the P7 mesentery, GATA2 and PROX1 expression was decreased in *Kras*^*+/G12D*^ mice compared to controls, but the expression levels of FOXC2 and VEGFR3 remained the same (Figure S3A-C). In the P21 diaphragm, PROX1 expression was decreased, while VEGFR3 expression was increased in the *Kras*^*+/G12D*^ mice. Additionally, FOXC2 expression was not upregulated to make valve territories in the P21 diaphragm of *Kras*^*+/G12D*^ mice (Figure S3D-E). In the P21 ear skin, the expression of FOXC2 and PROX1 was decreased, but VEGFR3 expression was increased in *Kras*^*+/G12D*^ mice when compared to controls (Figure S3F-G). Collectively, our data show that KRAS-G12D downregulates the transcription factors controlling valve development and leads to tissue-specific changes in their expression *in vivo*.

### The KRAS-G12D mutation does not alter VE-cadherin expression or localization *in vivo*

Two other studies on vascular malformations have shown that hyperactivation of the MAPK pathway leads to the loss of VE-cadherin from the cell membrane in cultured endothelial cells.^15,17^ Because VE-cadherin regulates mechanotransduction signaling in lymphatic vessels and is required for lymphatic valve formation and maintenance, we tested the hypothesis that the lack of VE-cadherin signaling is the cause of valve loss in *Kras*^*+/G12D*^ mice.^18^ To first confirm the previous results *in vitro*, we infected cultured hdLECs with a lentivirus expressing KRAS-G12D, or an identical vector in which KRAS-G12D was replaced by mCherry as a CV, and then exposed the cells to either static or OSS for 48 hours (Figure S2). The qRT-PCR and western blot results indicated that VE-cadherin (*CDH5*) was downregulated under both static and OSS conditions, at both the mRNA (Figure S2A) and protein levels (Figure S2B).

To evaluate the expression of VE-cadherin *in vivo*, we generated a reporter mouse where the monomeric red fluorescent protein (RFP), mScarlet, was fused to the cytoplasmic tail of VE-cadherin, which we named *Cdh5-mScarlet* (hereafter: *Cdh5*^*mS/mS*^). Whole-mount immunostaining was performed on *Flt4CreER*^*T2*^*;Kras*^*+/G12D*^*;Cdh5*^*+/mS*^*;Prox1GFP* mice and *Flt4CreER*^*T2*^*;Cdh5*^*+/mS*^*;Prox1GFP* controls using an anti-RFP antibody. Analysis of P14 mesenteries, P21 diaphragms, and P21 ear skin revealed the obvious presence of VE-cadherin at the cell membrane in all tissues analyzed from *Flt4CreER*^*T2*^*;Kras*^*+/G12D*^;*Cdh5*^*+/mS*^*;Prox1GFP* mice (Figure 3). Notably, the intensity of the staining was no different from the *Flt4CreER*^*T2*^*;Cdh5*^*+/mS*^*;Prox1GFP* controls. We further confirmed these results by performing whole-mount immunostaining using a monoclonal antibody specific for VE-cadherin which again revealed the presence of VE-cadherin at the cell membrane at similar intensities in the *Kras*^*+/G12D*^ and control mice (Figure S4). Thus, these data indicate that VE-cadherin is still present on the cell membrane in lymphatic vessels and therefore defective VE-cadherin-mediated mechanotransduction is not a likely explanation for the loss of lymphatic valves in *Kras*^*+/G12D*^ mice.

**Figure 3.**
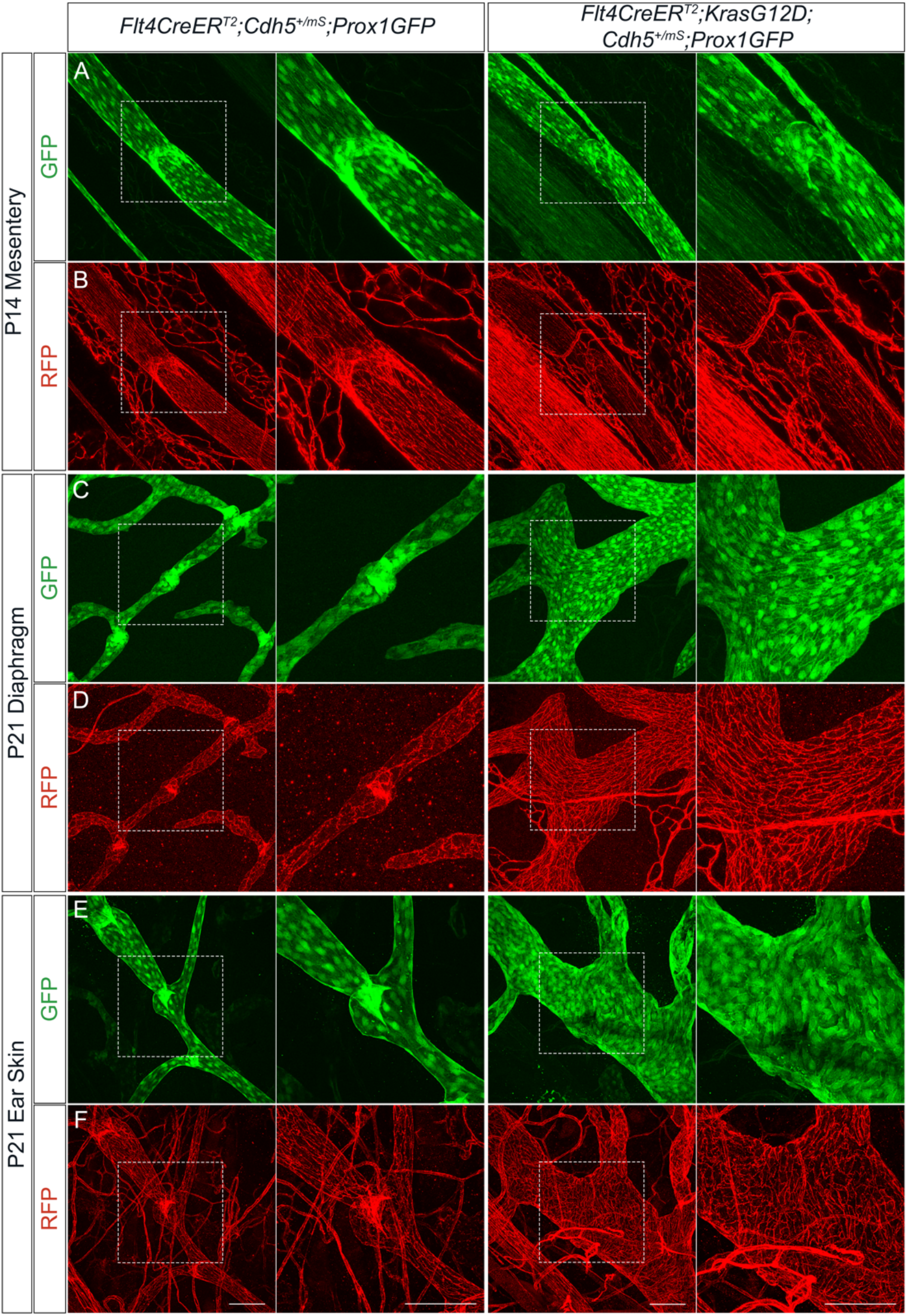
A fluorescent VE-cadherin fusion mouse reveals the presence of VE-cadherin in *Flt4CreER*^*T2*^*;Kras*^*+/G12D*^ tissues. *Cdh5-mScarlet* (*Cdh5*^*mS/mS*^) fusion mice were generated and bred with *Flt4CreER*^*T2*^*;Kras*^*+/G12D*^;*Prox1GFP* mice for evaluating the expression and localization of VE-cadherin (vascular endothelial-cadherin). Representative whole-mount immunostaining of postnatal day (P)14 mesenteries (**A**-**B**), P21 diaphragms (**C**-**D**), and P21 ear skin (**E**-**F**) from *Flt4CreER*^*T2*^ controls and *Flt4CreER*^*T2*^*;Kras*^*+/G12D*^ mice expressing the *Prox1GFP* reporter (GFP; green fluorescent protein; **A, C, E**) and *Cdh5*^*+/mS*^ (RFP; red fluorescent protein; **B, D, F**). Scale bars are 100 μm and n=3 for each tissue for **A** through **F**.

### Activating KRAS mutations lead to an upregulation of the PA pathway and MMPs in mice and hdLECs

We analyzed previously processed RNA-sequencing data from gene expression omnibus (GEO; GSE239737) of hdLECs that overexpressed KRAS-G12D or GFP as a control vector, which revealed the upregulation of several components of the PA pathway and MMPs, with MMP-9 being the most upregulated (Table 1).^30^

**Table 1.** Upregulation of the plasminogen activator (PA) pathway and matrix metalloproteinases (MMPs) in cultured lymphatic endothelial cells (LECs) expressing the KRAS-G12D mutation. Human dermal (hd) LECs were infected with a lentivirus expressing the KRAS-G12D mutation, or an identical vector expressing GFP as a control vector, and RNA-sequencing was performed.

| Gene | Protein | Log2 (Fold-Change) |
| --- | --- | --- |
| MMP-9 | Matrix Metalloproteinase-9 | 20.8 |
| MMP-1 | Matrix Metalloproteinase-1 | 7.20 |
| MMP-14 | Matrix Metalloproteinase-14 | 4.01 |
| MMP-10 | Matrix Metalloproteinase-10 | 3.02 |
| MMP-17 | Matrix Metalloproteinase-17 | 2.27 |
| MMP-2 | Matrix Metalloproteinase-2 | 0.46 |
| PLAT | Tissue plasminogen activator (tPA) | 6.77 |
| PLAU | Urokinase-type plasminogen activator (uPA) | 7.72 |
| PLAUR | Urokinase receptor | 6.81 |

To confirm these findings, we performed qRT-PCR on hdLECs infected with a lentivirus expressing the KRAS-G12D mutation, or an identical vector expressing wild type (WT) KRAS as an overexpression control, or an identical vector where KRAS was replaced by mCherry as a CV (Figure 4A-D). The results confirmed a significant upregulation of tPA (tissue plasminogen activator; *PLAT*), uPA (urokinase plasminogen activator; *PLAU*), uPAR (urokinase-type plasminogen activator receptor; *PLAUR*), and MMP-1 in the hdLECs expressing KRAS-G12D. The expression of MMP-2, MMP-9, MMP-10, and MMP-14 were also consistently upregulated but did not reach statistical significance due to the increased variance in the data, violating an assumption for one-way ANOVA. The expression of uPAR, MMP-1, and MMP-10 in hdLECs expressing WT KRAS was upregulated when compared to control hdLECs. PAI-1 (Plasminogen activator inhibitor 1; *SERPIN E1*) expression did not change. MMP-17 expression was downregulated in hdLECs expressing KRAS-G12D and in hdLECs expressing WT KRAS when compared to control hdLECs (Figure 4A). Western blot analysis confirmed an increase in uPAR and MMP-14 expression at the protein level in hdLECs expressing KRAS-G12D compared to CV or WT KRAS-expressing hdLECs (Figure 4B).

**Figure 4.**
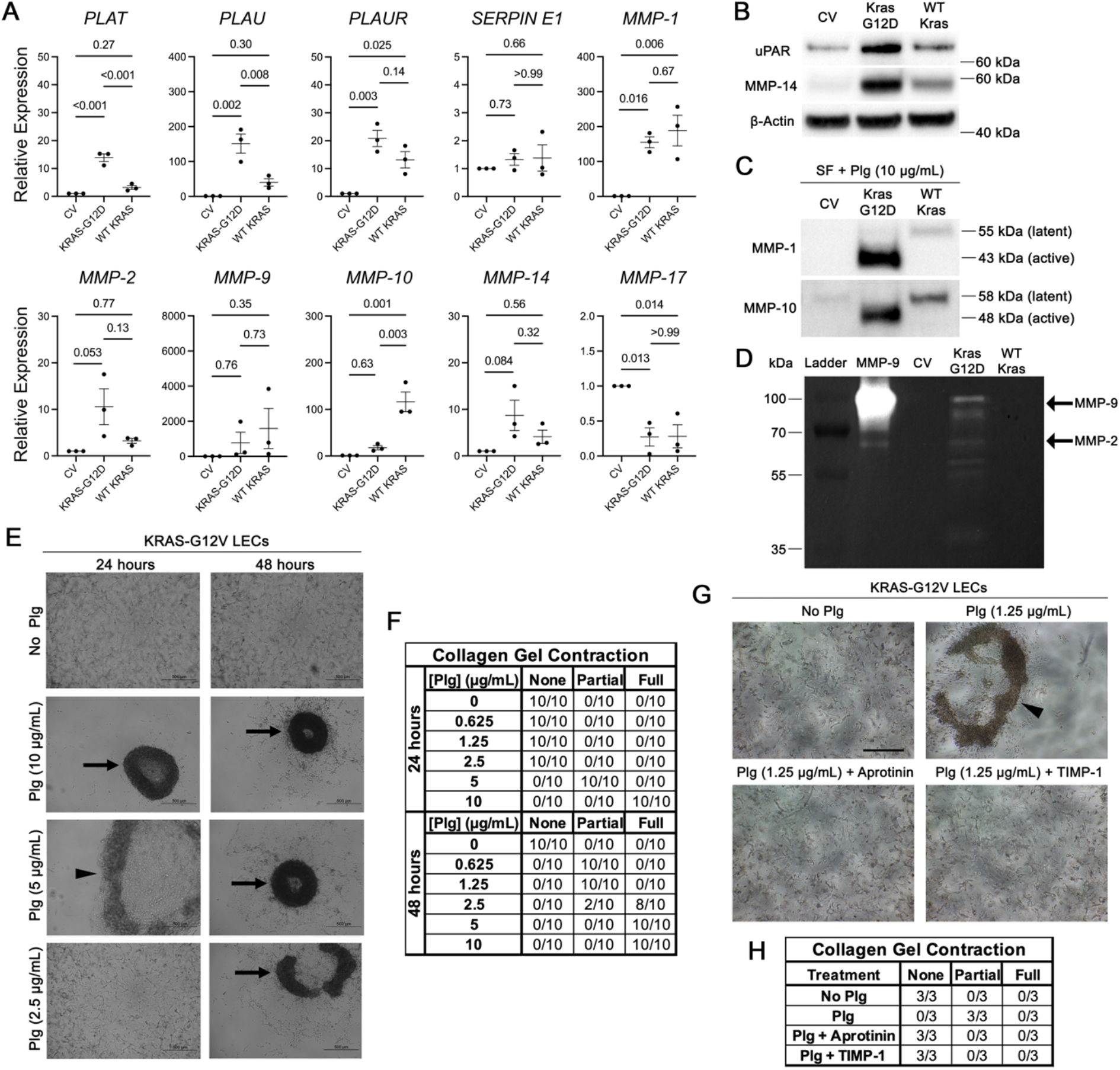
Upregulation of the plasminogen activator (PA) pathway and matrix metalloproteinases (MMPs) in cultured lymphatic endothelial cells (LECs) expressing activating KRAS mutations. **A**- **D**, Human dermal (hd) LECs were infected with a lentivirus expressing the KRAS-G12D mutation with a GFP (green fluorescent protein) reporter, or an identical vector expressing WT KRAS, or an identical vector where the KRAS was replaced by mCherry as a control vector (CV). **A**, qRT-PCR (quantitative real-time polymerase chain reaction) analysis of mRNA expression of the PA pathway and MMPs. All values are mean ± SEM of n=3. Analysis was performed with 1-way ANOVA with the Tukey multiple comparison test. **B**, Western blot analysis of uPAR (urokinase-type plasminogen activator receptor) and MMP-14 from protein lysates. β-actin was used as a loading control. **C**, Western blot analysis of MMP-1 and MMP-10 from conditioned media collected after hdLECs were incubated for 18 hours with serum-free (SF) media with added plasminogen (Plg). **D**, Gelatin zymography performed with conditioned media collected after hdLECs were incubated for 18 hours with serum-free (SF) media with added plasminogen (plg) indicates the presence of activated MMP-2 (64 kDa) and MMP-9 (82 kDa). **E**, hdLECs carrying KRAS-G12V were cultured in 3D collagen matrices under defined serum-free conditions in the absence or presence of the indicated concentrations of Plg in µg/ml. Cultures were photographed at either 24 or 48 hours of culture to evaluate the extent of collagen gel contraction, a measure of collagen type I matrix degradation. Arrows indicate fully contracted gels, and arrowheads indicate partially contracted gels. **F**, Quantification of the number of gels that were non-contracted, fully contracted or partially contracted at 24 or 48 hours (n=10). **G**, hdLECs carrying KRAS-G12V were cultured in 3D collagen matrices under defined serum-free conditions in the absence or presence of Plg at 1.25 µg/ml. The Plg cultures were left untreated or treated with 1 µg/ml of the serine protease inhibitor, aprotinin, or the MMP inhibitor, TIMP-1 (tissue inhibitor of metalloproteinases-1) at 5 µg/ml. Cells were photographed and quantitated (**H**) for gel contraction at 72 hours (n=3). Scale bar is 500 µm for **E** and **G**.

To assess the expression of secreted MMPs, we collected conditioned media from hdLECs that were incubated for 18 hours with serum-free (SF) media supplemented with plasminogen (Plg; Figure 4C-D). We added plasminogen to probe for the presence of tPA and uPA, as these enzymes cleave plasminogen into the active plasmin enzyme that can activate MMPs. We then performed western blot analysis for MMP-1 and MMP-10, which can distinguish the latent versus active forms of these MMPs by molecular weight. In the control hdLECs, no activated (43 kDa) or latent (55 kDa) MMP-1 were present. Similarly, no activated (48 kDa) MMP-10 was present and barely any latent (58 kDa) MMP-10 was present in control hdLECs. The hdLECs expressing KRAS-G12D had the highest expression of both MMP-1 and MMP-10 when compared to hdLECs overexpressing the CV or WT KRAS. The bands were of a lower molecular weight (MW), indicating that MMP-1 and MMP-10 were activated in KRAS-G12D hdLECs. In the hdLECs overexpressing WT KRAS, there was a slight upregulation of MMP-1 and MMP-10 compared to the control hdLECs, but both were the latent forms (Figure 4C). Gelatin zymography was performed with conditioned media using the same methods, and the results showed the presence of activated MMP-2 (64 kDa) and MMP-9 (82 kDa) only in hdLECs expressing KRAS-G12D (Figure 4D).

While the presence of activated MMPs suggest that hdLECs have the potential to degrade matrix proteins, like collagen, it does not provide direct evidence. To this end, hdLECs were infected with lentivirus expressing the activating KRAS-G12V mutation and cultured in a 3D matrix composed of collagen type I under defined serum-free conditions (Figure 4E-H). Plasminogen was added to the matrix in varying concentrations (0, 0.625, 1.25, 2.5, 5, 10 µg/ml). Cultures were photographed at 24 or 48 hours to evaluate the extent of collagen gel contraction, which is a measure of matrix degradation. In the presence of plasminogen, the KRAS-G12V hdLECs completely degraded the collagen matrix, causing its contraction into a ball. This process was accelerated with higher concentrations of plasminogen (Figure 4E). These results were quantified by counting the number of gels that were non-contracted, fully contracted, or partially contracted at 24 or 48 hours (Figure 4F). To determine if the collagen degradation could be prevented by inhibiting serine proteases or by inhibiting MMPs directly, hdLECs expressing KRAS-G12V were cultured in 3D collagen matrices under defined serum-free conditions in the absence or presence of plasminogen (1.25 µg/ml), and were then left untreated or treated with the broad-spectrum serine protease inhibitor, aprotinin (1 µg/ml), or the MMP inhibitor, TIMP-1 (tissue inhibitor of metalloproteinases-1; 5 µg/ml). Cultures were photographed 72 hours later, and results showed that aprotinin and TIMP-1 treatment both prevented the collagen gel contraction that occurs in the plasminogen-treated KRAS-G12V LEC cultures (Figure 4G), and this was quantified in Figure 4H.

To assess these results *in vivo*, we performed whole-mount immunostaining of membrane-bound uPAR, and two ECM proteins that are found in the core of the lymphatic valve, fibronectin and laminin α5, of P21 mesenteries, diaphragms, and ear skin of *Kras*^*+/G12D*^ mice and controls (Figure 5). Results showed upregulation of uPAR in the P21 mesentery (Figure 5A) and the P21 diaphragm (Figure 5D). Additionally, less fibronectin was found in the remaining lymphatic valve ECM core in the *Kras*^*+/G12D*^ mice when compared to controls in the P21 mesentery (Figure 5C). Less laminin α5 was present in both the valve and lymphangion LECs in *Kras*^*+/G12D*^ mice when compared to controls in the P21 mesentery (Figure 5B). However, in the P21 ear skin, less laminin α5 was present in the lymphangion LECs in *Kras*^*+/G12D*^ mice when compared to controls, but similar expression was observed in the remaining valve LECs (Figure 5E). Altogether, these data show that a main component of the PA pathway is upregulated *in vivo*, and that MMPs are not only upregulated, but also activated, leading to ECM protein degradation in the valve leaflet core.

**Figure 5.**
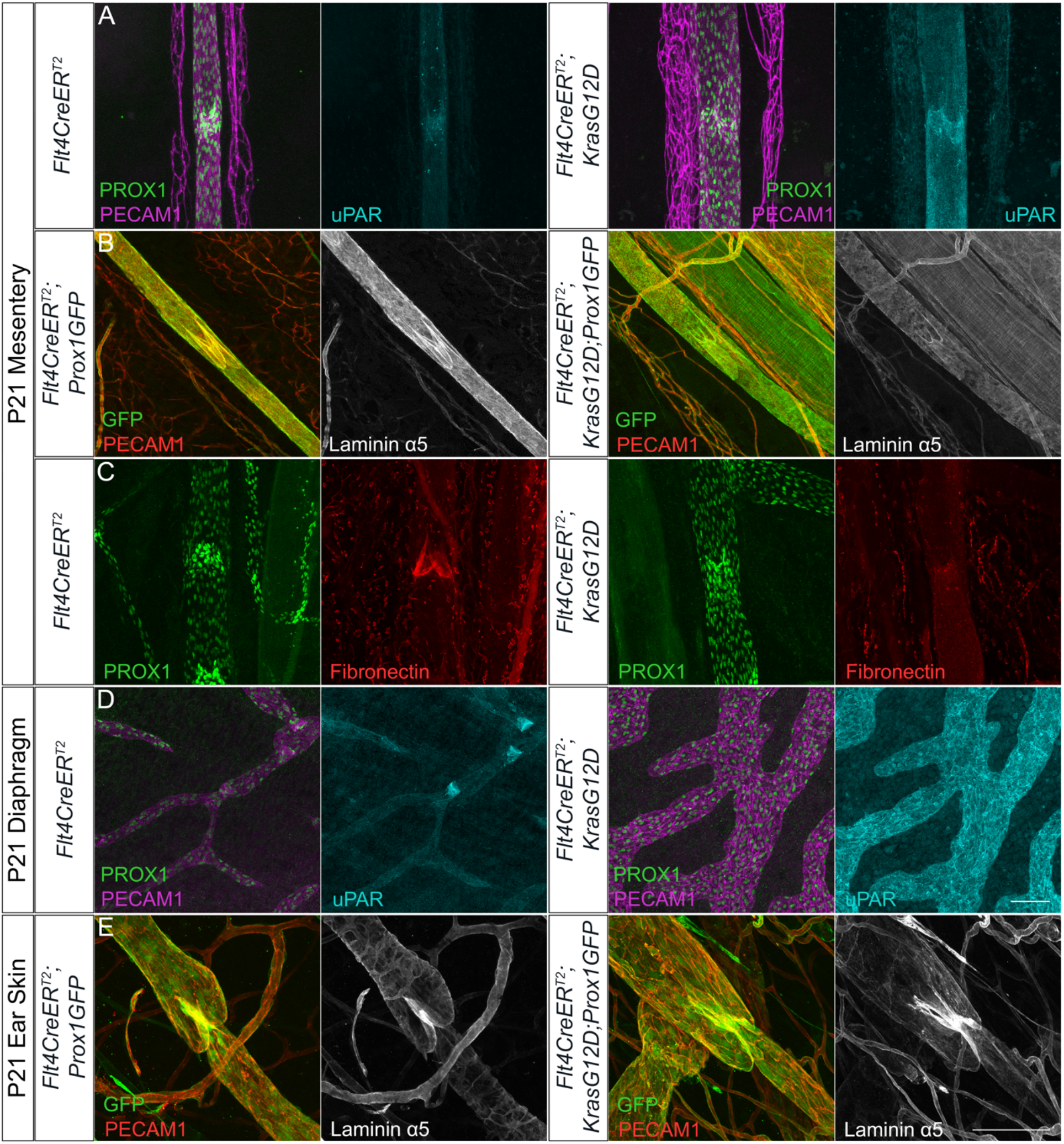
Plasminogen activator pathway and extracellular matrix proteins in *Flt4CreER*^*T2*^*;Kras*^*+/G12D*^ mice. Representative whole-mount immunostaining of postnatal day (P)21 mesenteries (**A**-**C**), P21 diaphragms (**D**), and P21 ear skins (**E**) from *Flt4CreER*^*T2*^ controls and *Flt4CreER*^*T2*^*;Kras*^*+/G12D*^ mice with PROX1 (prospero homeobox protein 1; green), PECAM1 (platelet and endothelial cell adhesion molecule 1; magenta), uPAR (urokinase-type plasminogen activator receptor; cyan; **A, D**), GFP (green fluorescent protein), PECAM1 (red), laminin α5 (white; **B, E**), PROX1 (green), and fibronectin (red; **C**). Scale bars are 100 μm and n=3 for each tissue for **A**-**E**.

### Plasminogen knockout does not rescue lymphatic valve defects in mice expressing the KRAS-G12D mutation

To determine if genetic ablation of plasminogen would rescue the lymphatic valve loss observed in *Kras*^*+/G12D*^ mice, we bred *Flt4CreER*^*T2*^*;Kras*^*+/G12D*^*;Prox1GFP* mice with global *Plg*^*-/-*^ mice that lack all plasminogen (Figure S5). Quantification of valves per millimeter (mm) of vessel at P21 in the mesentery, diaphragm, and ear skin revealed no significant difference between mice expressing *Kras*^*+/G12D*^ and those expressing both *Kras*^*+/G12D*^ and *Plg*^*-/-*^ (Figure S5D, S5H, S5L). Thus, removing plasminogen does not rescue lymphatic valve loss, suggesting that MMP activation persists downstream of the KRAS-G12D mutation, possibly via other serine proteases.

### Inhibiting MMPs rescues valve loss in the mesentery of mice expressing KRAS-G12D

Because MMPs can be activated by many serine proteases and other signaling pathways, we tested whether direct inhibition of MMPs was able to rescue the lack of lymphatic valves in the *Kras*^*+/G12D*^ mice. *Flt4CreER*^*T2*^*;Kras*^*+/G12D*^*;Prox1GFP* mice and *Flt4CreER*^*T2*^*;Prox1GFP* controls were administered ilomastat, a pan-MMP inhibitor, once every day from P0 until tissues were harvested at P7 or P14.

In the diaphragm at P7, the ilomastat-treated controls had a significant decrease in valves per millimeter when compared to vehicle-treated controls (0.46 valves/mm vs. 0.73 valves/mm, n=6; Figure S6A-E). As expected, the vehicle-treated *Kras*^*+/G12D*^ mice had a significant decrease in valves per millimeter when compared to vehicle-treated controls (0.12 valves/mm vs. 0.73 valves/mm, n=6). The ilomastat-treated *Kras*^*+/G12D*^ mice had a significant decrease in valves per millimeter when compared to vehicle-treated controls (0.09 valves/mm vs. 0.73 valves/mm, n=6), and did not differ from the vehicle-treated *Kras*^*+/G12D*^ mice (0.09 valves/mm vs. 0.12 valves/mm, n=6). In the diaphragm at P14, no significant difference in valves per millimeter between the ilomastat-treated and vehicle-treated controls was observed (0.68 valves/mm vs. 1.06 valves/mm, n=6; Figure S6F-J). As before, the vehicle-treated *Kras*^*+/G12D*^ mice had a significant decrease in valves per millimeter when compared to the vehicle-treated control mice (0.11 valves/mm vs. 1.06 valves/mm, n=6). The ilomastat-treated *Kras*^*+/G12D*^ mice had a significant decrease in valves per millimeter when compared to vehicle-treated control mice (0.09 valves/mm vs. 1.06 valves/mm, n=6), and did not differ from the vehicle-treated *Kras*^*+/G12D*^ mice (0.09 valves/mm vs. 0.11 valves/mm, n=6). Thus, ilomastat treatment did not rescue valve loss in the diaphragm at P7 or P14.

In contrast, in P7 mesenteries, ilomastat-treated controls and ilomastat-treated *Kras*^*+/G12D*^ mice exhibited a significant increase in valves per millimeter when compared to the vehicle-treated controls (1.48 valves/mm vs. 1.04 valves/mm, n=6) and vehicle-treated *Kras*^*+/G12D*^ mice (0.91 valves/mm vs. 0.42 valves/mm, n=6), respectively (Figure 6A-E). As expected, vehicle-treated *Kras*^*+/G12D*^ mice had a significant decrease in valves per millimeter when compared to vehicle-treated controls (0.42 valves/mm vs 1.04 valves/mm, n=6). Interestingly, no significant difference in valves per millimeter between the vehicle-treated control mice and the ilomastat-treated *Kras*^*+/G12D*^ mice was observed (1.04 valves/mm vs. 0.91 valves/mm, n=6), indicating that ilomastat treatment fully rescues valve loss in the mesentery at P7 (Figure 6A-E).

**Figure 6.**
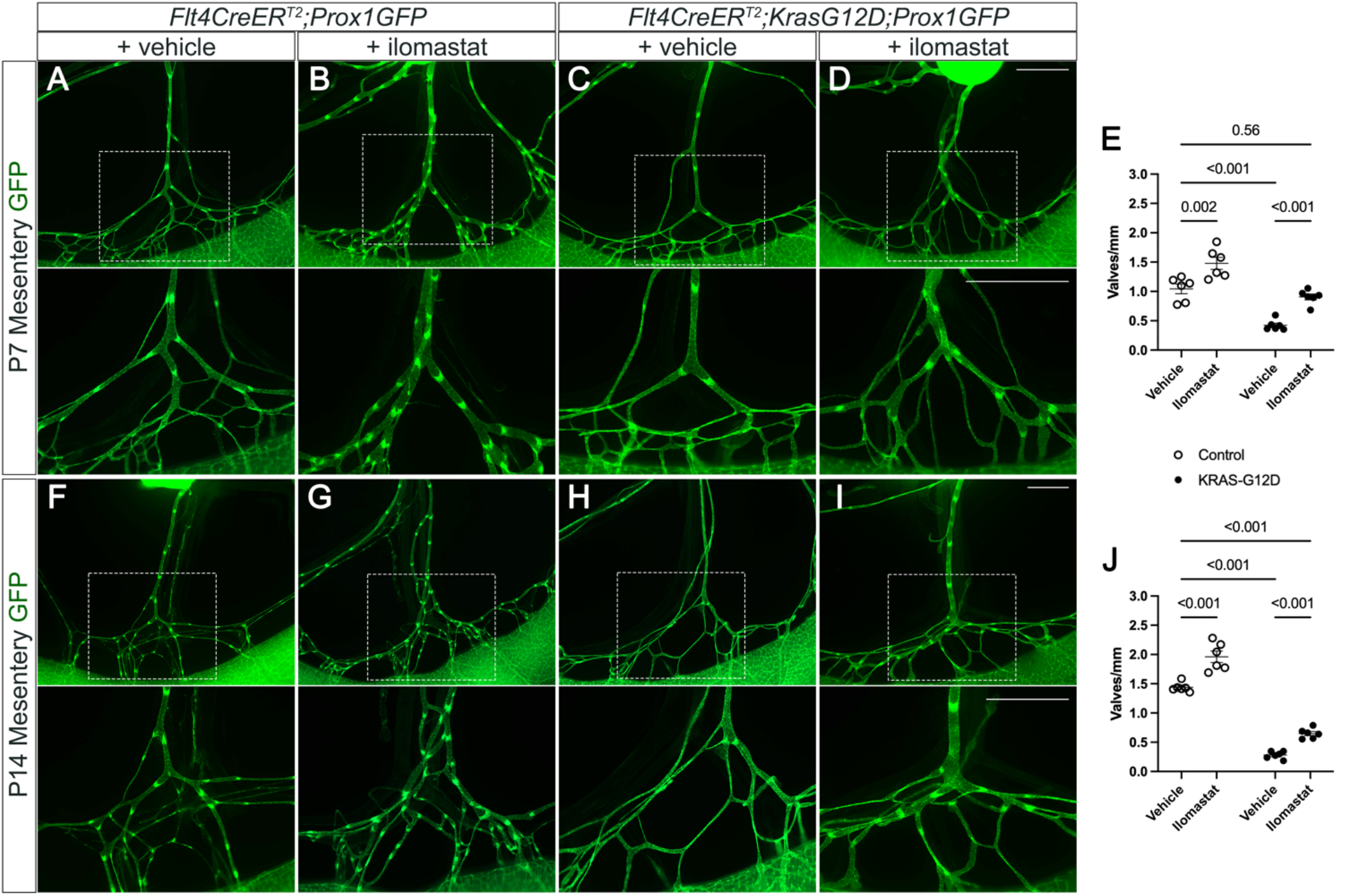
Ilomastat treatment rescues valve loss in the mesentery of *Flt4CreER*^*T2*^*;Kras*^*+/G12D*^ mice. Mice were treated with ilomastat, a pan-matrix metalloproteinase (MMP) inhibitor, every day from postnatal day (P)0 until tissues were harvested. Fluorescence images of the mesenteric lymphatic vasculature represented by green fluorescent protein (GFP) at P7 (**A**-**D**), and P14 (**F**-**I**). Scale bar is 1 mm. **E, J**, Valves per millimeter (mm) of vessel was quantified. All values are mean ± SEM of n=6 per genotype. Analysis was performed with 2-way ANOVA with the Tukey multiple comparison test.

In the P14 mesentery, ilomastat-treated controls had a significant increase in valves per millimeter when compared to the vehicle-treated controls (1.96 valves/mm vs. 1.44 valves/mm, n=6; Figure 6F-J). As expected, vehicle-treated *Kras*^*+/G12D*^ mice had a significant decrease in valves per millimeter when compared to vehicle-treated controls (0.28 valves/mm vs. 1.44 valves/mm, n=6). While ilomastat-treated *Kras*^*+/G12D*^ mice had a significant decrease in valves per millimeter when compared to vehicle-treated control mice (0.65 valves/mm vs. 1.44 valves/mm, n=6), we found a significant 2.3-fold increase in lymphatic valves compared to vehicle-treated *Kras*^*+/G12D*^ mice (0.65 valves/mm vs. 0.28 valves/mm, n=6). These data show that ilomastat treatment can rescue valve loss in the mesentery at P14 (Figure 6F-J).

## DISCUSSION

Recently, some patients who present with life-threatening symptoms such as chylothorax were found to carry activating KRAS mutations.^13^ Chylothorax and lymphatic valve loss were previously reported in a LM mouse model with an activating *Kras*^+/G12D^ mutation.^7^ However, the mechanism that causes lymphatic valve deficiency in mice expressing *Kras*^+/G12D^ is currently unknown. Here, we identify a putative mechanism for lymphatic valve loss in *Kras*^+/G12D^ mice. We found that although VE-cadherin is downregulated in cultured hdLECs expressing KRAS-G12D, these results are not recapitulated *in vivo*, and VE-cadherin remains present at the cell membrane in *Kras*^+/G12D^ lymphatic vessels. Instead, we show that activated KRAS signaling leads to an upregulation of the PA pathway and MMPs *in vitro* and *in vivo*. Further, we were able to rescue lymphatic valve loss in *Kras*^+/G12D^ lymphatic vessels by using the pan-MMP inhibitor, ilomastat.

Other studies have shown that KRAS-G12D expressed by LECs leads to lymphatic valve deficiencies in the mesentery, ear skin, and lungs,^7,8^ and we also show that the loss of lymphatic valves occurs in multiple organs throughout the body at different timepoints. Our data additionally provide evidence that KRAS-G12D prevents the formation of valves because lymphatic vessels grow into the diaphragm postnatally (data not shown) and newborn pups lack ears, which grow completely postnatally (Figure 2). A previous report established a correlation between chylothorax and valve loss in mice,^6^ and results from this study and others further support that chylothorax caused by lymphatic KRAS-G12D is due to the lack of valves (Figure 1).^7,8^

Activating mutations in the MAPK pathway have been shown to cause a loss of VE-cadherin localization at the cell membrane *in vitro*.^15,17^ Our results confirm that VE-cadherin is downregulated on the RNA and protein levels in cultured hdLECs expressing an activating KRAS-G12D mutation (Figure S2). We previously demonstrated that VE-cadherin regulates mechanotransduction signaling in lymphatic vessels, and genetic knockout of VE-cadherin leads to a similarly severe loss of lymphatic valves as *Kras*^*+/G12D*^.^18^ Thus, if VE-cadherin was downregulated, degraded, or internalized *in vivo*, it logically follows that mechanotransduction signaling would be inhibited in lymphatic endothelium and would explain the lack of valves. Upon assessing VE-cadherin expression *in vivo*, our data instead demonstrated that VE-cadherin is indeed present at the cell membrane in lymphatic vessels expressing KRAS-G12D, with no noticeable difference in the expression level (Figure 3, S4). Therefore, VE-cadherin is not downregulated in lymphatic vessels expressing the KRAS-G12D mutation. Future studies should assess VE-cadherin expression in blood vascular malformations and LMs expressing other MAPK gene mutations.

The PA pathway is generally known for its role in fibrinolysis, which is how fibrin blood clots are dissolved.^31,32^ Plasminogen is synthesized by the liver and then secreted into the bloodstream where it is cleaved into plasmin by two serine proteases, tPA and uPA.^32^ uPA must bind to uPAR to become activated.^33^ Plasmin is a serine protease that can cleave ECM proteins and activate the pro-peptide forms of matrix metalloproteases (MMPs).^34,35^ To safeguard against excessive fibrinolysis, two other proteins inhibit tPA/uPA activity, plasminogen activator inhibitor 1 (PAI-1) and plasminogen activator inhibitor 2 (PAI-2).^33^ PAI-2 is not expressed in LECs, as found in RNA-sequencing data published from two previous studies.^36,37^ We found that multiple proteins in the PA pathway and several MMPs were upregulated in cultured hdLECs expressing KRAS-G12D (Table 1, Figure 4). Interestingly, these upregulated MMPs degrade the same ECM proteins found in the valve leaflet core (e.g., laminin-α5, collagen, and fibronectin-EIIIA).^2,3,38^ This led us to hypothesize that hyperactive KRAS signaling increases the expression of MMPs and key PA enzymes that then activate the upregulated MMPs to degrade the ECM of lymphatic valves. Based on a previous study and published RNA-sequencing data, there are a number of other ECM proteins (such as tenascin-C, nidogen, and perlecan) expressed in the valve leaflet that can be degraded by these same MMPs.^2,36,38^ Our hypothesis is supported by western blot showing that MMP-1 and MMP-10 were upregulated and activated only in the hdLECs expressing the KRAS-G12D mutation.

Additionally, our data show that hdLECs expressing KRAS-G12D have the capability to convert plasminogen into plasmin, and have elevated message for the tPA and uPA enzymes. *In vivo*, more plasminogen would be cleaved into plasmin that could then activate MMP-1 and MMP-10. In gel zymography, we found that MMP-2 and MMP-9 are expressed in the hdLECs expressing KRAS-G12D, but not in hdLECs expressing CV or WT KRAS. Lastly, our data show that PAI-1 (*SERPIN E1*) expression does not change in cultured hdLECs expressing KRAS-G12D, indicating that the upregulation of the PA pathway is not inhibited by PAI-1 (Figure 4). These results are supported by published literature showing that the MAPK pathway regulates MMP expression.^39-45^ hdLECs expressing an activating KRAS-G12V mutation cultured in 3D collagen matrices were treated with plasminogen which led to contraction and degradation of the collagen gel, which was prevented by adding TIMP-1 (MMP inhibitor) or aprotinin (serine protease inhibitor). While this experiment shows that plasmin is sufficient for MMP activation and collagen degradation, it does not test whether plasmin is required. Global knockout of plasminogen in the *Kras*^*+/G12D*^ mice failed to rescue lymphatic valve loss, therefore revealing that plasmin is not required for MMP activation and that any plasmin generated is not playing a significant role in valve ECM core degradation. However, multiple other pathways can activate the upregulated MMPs. The activation of MMP-14 is governed by proprotein convertases such as furin,^46^ and furin is expressed in LECs,^36^ and upregulated by the KRAS-G12D mutation in the RNA-sequencing data (GSE239737).^30^ KLK14 (kallikrein-related peptidase 14) has also been shown to activate MMP-14.^47^ Other serine proteases such as plasma kallikrein, trypsin, neutrophil elastase, cathepsin G, tryptase, and chymase can activate MMP-1 and MMP-10.^48^ Increased levels of MMP-10 can activate MMP-1.^48^ MMP-10 can be activated by auto-activation or exposure to free-radicals.^48^ KLK7 (kallikrein-related peptidase 7) was previously shown to activate MMP-9.^49^ uPA itself can activate MMP-9,^50^ and tPA can lead to increased MMP-2 and MMP-9 expression, and can also activate MMP-9.^51^ Hepsin is a type II transmembrane serine protease that has been shown to activate MMPs.^52^ All of these are alternate ways in which MMPs can be activated without plasminogen/plasmin.

Lymphatic valve defects in the *Kras*^*+/G12D*^ mice were fully rescued by ilomastat in the mesentery at P7 and significantly rescued at P14, identifying a role for active MMPs in lymphatic valve degradation. Activating RAS/MAPK mutations have been identified in arteriovenous malformations (AVMs),^53^ and treatment of an AVM in the right upper extremity with the broad-spectrum MMP inhibitor, marimastat, led to extreme pain relief, improvement in mobility, and regression of bone erosion and soft tissue swelling^54^. Together with our results, this supports a role for MMPs in the pathology of vascular malformations caused by hyperactivation of the MAPK pathway. Multiple reports indicate that RAS activation leads to basement membrane and ECM protein degradation.^25,55,56^ One study showed that excessive MAPK activation mediated by KRAS-G12D leads to lymphangiectasia through the disruption of lymphatic basement membrane development and composition, thus promoting chylous effusion in the pleural space. Chylous effusion then inhibits gas exchange in alveoli and can lead to neonatal death^55^. A separate report expressed KRAS-G12V in an *in vitro* model allowing human BECs to undergo tubulogenesis. Expression of KRAS-G12V caused decreased basement membrane deposition and an increase in dilated vessels, and this was partly due to overactive MT1-MMP (MMP-14), which degrades basement membrane proteins^56^. The same authors performed another study, using the same *in vitro* model showing that KRAS-G12V-expressing BEC tubes have a higher likelihood to regress due to enhanced serine protease-mediated activation of MMP-1.^25^ Collectively, these studies support our hypothesis that MAPK hyperactivation leads to ECM degradation in lymphatic valve leaflets.

While we provide new insights into the pathobiology of lymphatic malformations with activating KRAS mutations, our study has a few limitations. We could not assess the *in vivo* expression of uPA, tPA, or MMPs because these enzymes are secreted from cells and collecting small amounts of lymph is technically challenging. Another limitation of this study is that although ilomastat is a pan-MMP inhibitor, it does not inhibit all MMPs. More specifically, it inhibits all the upregulated MMPs in this study except for MMP-10, and inhibits each MMP at a different potency (based on the K_i_ values). Lastly, our rescue of lymphatic valve loss with ilomastat treatment was tissue-dependent, which may indicate that different tissues have different mechanisms of valve loss (e.g. dependency on MMP-10). Alternatively, drug bioavailability and tissue distribution are two other factors that could explain why we observed a tissue-specific effect of ilomastat treatment on lymphatic valve growth and regression.

Even though rapamycin and trametinib have been used for treating LMs, they have significant side effects that may reduce patient compliance, and neither of these drugs directly inhibit the downstream targets that cause the loss of lymphatic valves. Our study identifies a novel pathway responsible for impaired lymphatic valve development, which can be pharmacologically targeted, and may lead to novel combinatorial treatments for LMs by inhibiting the downstream targets that directly cause valve loss.

## Non-standard Abbreviations and Acronyms

AVM: arteriovenous malformation
BEC: blood endothelial cell
CV: control vector
ECM: extracellular matrix
FOXC2: forkhead transcription factor C2
GATA2: GATA-binding factor 2
hdLEC: human dermal lymphatic endothelial cell
KRAS: Kirsten rat sarcoma viral oncogene homolog
LEC: lymphatic endothelial cell
LM: lymphatic malformation
MAPK: mitogen-activated protein kinase
MMP: matrix metalloproteinase

## Acknowledgements

We thank Dr. Young-Kwon Hong for kindly providing the Prox1GFP strain. We thank Dr. Lydia Sorokin for kindly gifting the laminin-α5 antibody.

## Sources of Funding

This study was supported by National Institutes of Health grants R01 HL164825 and R01 HL142905 to J.P. Scallan, R01 HL145397 and R01 HL16698 to Y. Yang, T32 HL160529 and F31 HL174150 to D.M. Mastrogiacomo, Department of Defense grants W81XWH2110653 to J.P. Scallan, W81XWH2110652 to M.T Dellinger, and American Heart Association grant 24PRE1195095 to D.M. Mastrogiacomo.

## Disclosures

None.

## Supplemental Material

Supplemental Methods

Figure S1–S6

Major Resources Table

## Novelty and Significance

### What is Known?

- Multiple activating gene mutations in the MAPK pathway (ARAF, BRAF, KRAS, etc.) were recently identified in lymphatic malformations (LMs) that are associated with lymphedema, chylous ascites, and chylothorax.
- Cultured blood and lymphatic endothelial cells (LECs) expressing activating mutations in the MAPK pathway demonstrate a loss of VE-cadherin from the cell surface, but VE-cadherin has not been investigated *in vivo*.
- Lymphatic-specific expression of KRAS-G12D in a mouse model of LMs leads to a severe reduction in lymphatic valves, but the mechanisms responsible remain unknown.

### What New Information Does This Article Contribute?

- Lymphatic-specific expression of KRAS-G12D leads to upregulation of the plasminogen activator (PA) pathway and MMPs.
- VE-cadherin expression is unaltered *in vivo* in lymphatic vessels expressing KRAS-G12D, and therefore does not underlie the pathogenesis of LMs.
- Inhibition of MMPs led to a rescue of lymphatic valves in a mouse model of LMs expressing KRAS-G12D, and could be a potential new therapeutic target for combinatorial treatment.

Our studies reveal a mechanism of lymphatic valve loss in LMs caused by activating KRAS mutations. Hyperactivation of the MAPK pathway leads to an upregulation of the PA pathway and MMPs, thus causing degradation of the ECM core of developing and existing lymphatic valves, explaining the observed deficiency of valves in LM models. The current pharmacological treatments for LMs exhibit toxic side effects and do not directly inhibit the downstream proteins that degrade lymphatic valves. Our study provides insights into the pathophysiology of LMs and identifies potential combinatorial treatments for LMs that cause life-threatening chylothorax.

## Notes

### Competing Interest Statement

The authors have declared no competing interest.

